# ursaPGx: a new R package to annotate pharmacogenetic star alleles using phased whole genome sequencing data

**DOI:** 10.1101/2023.07.24.550372

**Authors:** Gennaro Calendo, Dara Kusic, Jozef Madzo, Neda Gharani, Laura Scheinfeldt

## Abstract

Long-read sequencing technologies offer new opportunities to generate high confidence phased whole genome sequencing data for robust pharmacogenetic annotation. Here we describe a new user-friendly R package, ursaPGx, designed to accept multi-sample phased whole genome sequencing data VCF input files and output star allele annotations for pharmacogenes annotated in PharmVar.

## Background

Pharmacogenomics (PGx) benefits medication management [1-7], however, pharmacogenetic annotation is often quite complex. Functional PGx annotation and corresponding clinical PGx recommendations rely on star (*) allele annotation [8, 9]; star alleles are often defined by more than one genetic variant [10-12]; and when the star allele defining variants are heterozygous, phased haplotype information is needed to resolve the annotation. In addition, annotations may change over time as new variants are characterized and incorporated into clinical PGx recommendations. Many resources and off the shelf tools are available to support researchers and clinicians interested in PGx annotation. Several tools are well suited for PGx annotation of unphased data (e.g., StellarPGx, Stargazer [13, 14]), and tools such as PharmCAT, while not computationally streamlined for multi-sample annotation, go a step further to incorporate clinical recommendations into the software output [15].

New long-read sequencing technologies offer opportunities to generate high confidence phased whole genome sequencing (WGS) data for robust PGx annotation. Here we describe ursaPGx, an R package designed to complement existing tools that leverages phased whole genome sequencing data for PGx annotation. ursaPGx is designed to run on a typical laptop computer using multi-sample, phased, WGS VCF files and provides an output table of star allele annotations for selected pharmacogenes annotated in PharmVar.

## Materials and Methods

### Samples

Phased multi-sample VCF files were downloaded for each of the star allele containing chromosomes from the 1000 Genomes Project. These VCF files were generated by the New York Genome Center for 3,202 1000 Genomes Project samples by aligning the 30x WGS reads to GRCh38 and performing SNV and INDEL variant calling as described in [16].

### Benchmark data

The accuracy of the star allele calling algorithm of ursaPGx was benchmarked against the next generation sequencing consensus calls generated by the Genetic Reference and Testing Material Coordination Program (GeT-RM) for *CYP2C8, CYP2C9*, and *CYP2C19* which combined the output of Astrolabe, Stargazer, and Aldy across investigator groups to generate a uniform diplotype call for each of the 137 samples included in their study [17], of which 87 also have 30x WGS data [16]. *CYP2D6* calls generated by ursaPGx’s implementation of Cyrius were benchmarked against calls generated by Chen et al. [18].

### Implementation and algorithm description

Users may choose any phased WGS VCF file of interest for use as input to ursaPGx. ursaPGx assigns phased diplotype calls from single-sample or multi-sample indexed VCF files using publicly available star allele definitions from PharmVar [10-12]. First, for a given pharmacogene, star allele defining positions are used to extract genotype data for all samples in the VCF. Next, the extracted positions are checked against each PharmVar haplotype definition in order to determine ‘callable’ alleles. In this context, a callable allele is defined as a haplotype definition where all allele defining variants are present in the sample VCF. Downstream analysis is then limited to the set of callable alleles. The set of callable alleles is then used to generate a genomic position by haplotype definition reference matrix. The cells of the reference matrix contain the nucleotide which defines the given haplotype for each of the positions present in the sample VCF. Positions that are not part of a given haplotype definition are filled with the reference nucleotide for the position. Using this reference matrix allows ursaPGx to disambiguate star allele definitions such as *CYP2C19*^*\**^*2* and *CYP2C19*^*\**^*35*, which share the same core allele definitions (CYP2C19*2, non-reference alleles for rs4244285, rs12769205, rs3758581; CYP2C19*35, non-reference alleles for rs12769205, rs3758581) and therefore must be distinguished by using a SNV unique to *CYP2C19*^*\**^*2* (rs4244285). After constructing the reference matrix, genotype calls are converted to their nucleotide representation and split into haplotype strings for each sample. For each sample, each haplotype string is checked for exact matches against all columns of the reference matrix. All exact matches to the reference for each sample haplotype string are reported for each sample. If no exact matches occur, then the haplotype call for that sample is reported as ambiguous (*Amb). Haplotype calls for each sample are then combined to form a single diplotype call for the given pharmacogene for each sample included in the VCF.

*CYP2D6* star allele calling in ursaPGx is performed with a modified version of Illumina *CYP2D6* star allele caller Cyrius, designed to function in R. The *CYP2D6* haplotype calling algorithm implemented in Cyrius is fully described in [18]. Briefly, Cyrius uses WGS BAM files to estimate the total number of copies of *CYP2D6* and *CYP2D7*, determines the number of complete *CYP2D6* and hybrid genes and uses these to estimate SVs impacting *CYP2D6* annotation. Cyrius then performs small variant calling for star allele defining positions and derives an estimate of their copy number, and then matches these calls and SVs against star allele definitions from PharmVar (7/15/2020) in order to produce final diplotype calls for each sample.

### Software

ursaPGx is a freely available and open source package implemented in the R programming language [19] and utilizes the *VariantAnnotation* package [20] from the Bioconductor project to provide a consistent interface with existing R packages for the analysis of genetic variant data. Star allele definitions in VCF format are downloaded from PharmVar (current version 5.2.13) and parsed into R objects. All package code and analysis scripts are hosted on GitHub (https://github.com/coriell-research/ursaPGx).

### Requirements

ursaPGx is designed to run on a personal laptop. Star allele calling for all 3,202 1000 Genomes Project samples for all 12 pharmacogenes takes ∼45 seconds on a 3.7 GHz 6-Core Intel Core i5 iMac. Cyrius *CYP2D6* calling implemented in ursaPGx takes ∼4 seconds per sample BAM.

## Results

*CYP2C8, CYP2C9*, and *CYP2C19* concordance was assessed for samples with matching IDs from the 30x WGS data in the GeT-RM benchmarking data sets (87/137) [17]. *CYP2D6* concordance was tested against diplotype calls from Chen et al. [18] in order to ensure accuracy of the Cyrius implementation within ursaPGx. Diplotype calls produced by ursaPGx were found to be highly consistent with those generated by GeT-RM for all four benchmarked pharmacogenes (**Table 1, Table S1**). For the 87 samples with matching IDs between the 1000 Genomes Project 30x WGS data and the GeT-RM NGS consensus benchmarking data, *CYP2C8* was found to be perfectly concordant. For *CYP2C19*, one subject sample (NA19122) was reported as *2|*Amb according to ursaPGx whereas the GeT-RM consensus call for this sample was reported as *2/*35. In the phased 30x WGS dataset, one haplotype was an exact match for *CYP2C19*2* but the other haplotype had no exact match to any PharmVar definition. Assuming accurate phasing of the input 30x WGS dataset, ursaPGx reports the inexact match as ambiguous for this sample.

**Table 1.** Concordance of ursaPGx diplotype calls with benchmarking datasets.

| **Gene** | **Concordance** | **Benchmarking Data** |
| --- | --- | --- |
| *CYP2C8* | 1.00 (87/87) | GeT-RM [17] |
| *CYP2C9* | 0.97 (84/87) | GeT-RM [17] |
| *CYP2C19* | 0.99 (86/87) | GeT-RM [17] |
| *CYP2D6* | 0.99 (2502/2504) | Cyrius [18] |

For *CYP2C9*, three samples were found to be discordant between ursaPGx and GeT-RM reported consensus calls. Two of the subject samples with discordant *CYP2C9* calls, NA19143 and NA19213, were annotated as *1/*6 by GeT-RM whereas ursaPGx assigned these samples as *1|*1. Because the *CYP2C9*6* defining variant (rs9332131) is not present in the phased 30x WGS dataset, *CYP2C9*6* is not included as a callable allele by ursaPGx and thus is not reported for these samples. One subject sample, HG01190, was assigned as *61|*1 by ursaPGx whereas GeT-RM reported the diplotype as *2/*61. However, this sample was found to be inconsistently annotated across labs in the GeT-RM benchmarking data with a minority subset of three of the annotation approaches assigning ^*^1/^*^61. Additionally, in the 30x WGS dataset, rs1799853 and rs202201137 are both heterozygous, and the non-reference allele for rs1799853 (*CYP2C9*2*) is on the same phased chromosome as the rs202201137 non-reference allele (presence of both non-reference alleles on the same haplotype defines the *61 variant according to PharmVar). Given the phase information from the 30x WGS, *61|*1 is the diplotype most consistent with the observed data for this sample.

Since Cyrius has already been shown to produce highly accurate *CYP2D6* star allele calls [18], we benchmarked ursaPGx’s implementation of Cyrius against the 2,504 Phase 3 1000 Genomes Project samples analyzed in the Cyrius publication in order to ensure that changes made to Cyrius, which were needed to port the software to R, were consistent with the original Cyrius implementation. 2,502 of the 2,504 samples were found to be exact matches with the Cyrius reported results. For the two discordant samples, NA18611 and HG02490, ursaPGx reported diplotype calls for these samples (*10/*2 and *2/*33, respectively) whereas the Cyrius benchmark did not assign a diplotype for these samples. This discrepancy is likely due to differences in BAM file input and downstream processing used in the 1000 Genomes Project NYGC 30x WGS data versus the WGS dataset used in the Cyrius publication [18].

## Discussion

Here we describe a new pharmacogenetic annotation tool, ursaPGx, that is designed to complement existing tools by leveraging multi-sample phased WGS data and PharmVar annotations. ursaPGx is implemented as an efficient and user-friendly R package that provides a simple interface for assigning star allele diplotypes to samples for PharmVar annotated genes including *CYP2D6*, by integrating the Cyrius *CYP2D6* star allele caller. ursaPGx is especially well suited to long-read WGS datasets (e.g., PacBio HiFi) where phasing confidence is high.

Our benchmarking analysis demonstrated high concordance, 100%, 97% and 99%, respectively for the three overlapping pharmacogenes, *CYP2C8, CYP2C9*, and *CYP2C19* included in the most recent GeT-RM report [17]. Two of the discordant samples for *CYP2C9* result from a star allele defining variant (^*^6) that is present in the GeT-RM dataset but not occurring in the 30x WGS 1000 Genomes Project dataset used to benchmark ursaPGx. The third discordant *CYP2C9* sample (HG01190) results presumably from differences in phasing and variant calling results. Finally, as detailed in the methods section above, when no perfect match to any PharmVar defined haplotype occurs, the ursaPGx output will be ‘*Amb’, and this implementation approach explains the single discordant *CYP2C19* sample, NA19122.

As with any annotation approach, ursaPGx includes several limitations. First and foremost, any error or missing variants in the input VCF file will propagate into errors in annotation. Similarly, any errors or uncertainty in phase will propagate into annotation errors, particularly when heterozygotes are phased incorrectly. In addition, our annotation approach is limited to the pharmacogenes annotated in PharmVar [10-12] and requires already phased input data. This annotation choice is specifically designed to take advantage of increasingly common long-read WGS datasets, such as the data being generated by the Human Pangenome Reference Consortium [21].

## Conclusion

New long-read sequencing technologies offer opportunities to generate high confidence phased whole genome sequencing data for robust PGx annotation. Here we describe ursaPGx, a user-friendly R package that leverages multi-sample phased whole genome sequencing data for star allele annotation.

## Availability and requirements

Project name: ursaPGx

Project home page: https://github.com/coriell-research/ursaPGx

Operating system(s): Platform independent

Programming language: R

Other requirements: None

License: GC will check Cyrius

Any restrictions to use by non-academics: GC will check Cyrius

## Declarations

### Ethics approval and consent to participate

All human data used in this study is publicly available through the 1000 Genomes Project [16].

### Availability of data and materials

NYGC WGS data (VCF files) are available through the www.internationalgenome.org website (https://www.internationalgenome.org/data-portal/data-collection/30x-grch38), and can be accessed from the following website link (https://ftp.1000genomes.ebi.ac.uk/vol1/ftp/data_collections/1000G_2504_high_coverage/working/20220422_3202_phased_SNV_INDEL_SV/). The version we used for the current study were last modified on 2022-11-14 08:33.

All package code and analysis scripts are hosted on GitHub (https://github.com/coriell-research/ursaPGx).

## Author’s Contributions

GC designed and implemented the project and contributed to writing and editing the manuscript; LS designed the project and contributed to writing and editing the manuscript. DK, JM, NG contributed to the design of the project, testing the software, and contributed to writing and editing the manuscript.

